# Microporous annealed particle scaffolds avoid foreign body response by down regulating complement-fibroblast-macrophage signaling loop

**DOI:** 10.64898/2026.04.17.719225

**Authors:** Colleen A. Roosa, Ethan Nicklow, Jeremy Ortmann, Riley Hannan, Jeffrey M. Sturek, Daniel Abebayehu, Donald R. Griffin

**Affiliations:** Department of Biomedical Engineering, University of Virginia, Charlottesville, VA; Department of Chemical Engineering, University of Virginia, Charlottesville, VA; Department of Medicine, Division of Pulmonary and Critical Care, University of Virginia, Charlottesville, VA

**Keywords:** Biomaterials, Foreign Body Response, Scaffold Porosity, Complement Pathway, Immune Modulation

## Abstract

Biomaterial implantation can trigger a foreign body response (FBR) that impedes tissue-implant integration. To investigate how implant porosity influences this response, we compared the immune response to subcutaneous implants of microporous annealed particle (MAP) scaffolds and nanoporous hydrogels using mass cytometry, single-cell RNA sequencing, and multiplex cytokine assays. MAP scaffolds promoted vascularization and tissue integration, marked by increased endothelial and regulatory T cells, and reduced proinflammatory immune cells and cytokines. In contrast, nanoporous hydrogels demonstrated enrichment of basophils, natural killer cells, and macrophage populations associated with fibrosis. Transcriptomic and proteomic analyses revealed that MAP scaffolds suppressed activation of the complement-fibroblast-macrophage signaling loop, particularly the C5a signaling crosstalk pathway. This was confirmed using C5-deficient mice, where complement-driven cytokine production was significantly reduced only in nanoporous implants. These findings demonstrate that scaffold porosity modulates immune and complement responses, identifying a key mechanism by which MAP scaffolds reduce FBR and improve biomaterial integration.

## INTRODUCTION

Biomaterial implantation initiates a wound environment that activates the host immune response.^1,2^ If the host inflammatory response is not resolved, a more chronic form of inflammation known as the foreign body response (FBR) can occur, which can lead to fibrous encapsulation of the implant that impairs integration and long-term function.^3^ Strategies such as surface modification^4–6^, attachment of bioactive molecules^7,8^, and release of anti-inflammatory drugs^9,10^ have been used to reduce the FBR to biomaterial implants. While these approaches can be effective in the short term, each faces long-term limitations: surface passivation can reduce tissue integration, bioactive molecules have limited half-lives, and drug-delivery systems eventually deplete their payload. An alternative approach, supported by the work of Ratner et. al, has shown that biomaterials with cell-scale porosity can reduce the FBR and increase pro-resolutory macrophage polarization.^11,12^ Additionally, we have found that the implantation of poly(ethylene) glycol (PEG) nanoporous (NP) hydrogels leads to the formation of a fibrotic capsule, which is due, in part, to insufficient implant porosity^13^. Using our MAP hydrogel scaffold with a chemically identical formulation to the NP hydrogel, we found that an open pore structure is sufficient to avoid observable scaffold-directed FBR^14^. This includes a minimal baseline immune response, negligible fibrotic encapsulation, and extensive cell infiltration. While we have observed a lack of FBR to MAP scaffolds using histological analysis, investigation into the transcriptomic and proteomic differences that influence the host response to NP and MAP hydrogel implants would offer greater mechanistic insight.

FBR is triggered by protein adsorption onto the biomaterial surface, followed by the adhesion of neutrophils, monocytes, and macrophages to the material.^15^ Subsequently, macrophages fuse together to form foreign body giant cells (FBGCs), which are a hallmark of the foreign body response.^2^ These FBGCs recruit and communicate with fibroblasts that localize to the implant surface and deposit layers of collagen and other extracellular matrix proteins that lead to a fibrotic encapsulation. In addition to protein and cell localization to the biomaterial surface, the FBR is also canonically guided by the activation of the complement pathway, which is a group of sequentially reacting proteins that mediate the host immune response to foreign pathogens and objects.^16^ Specifically, the complement system is involved in distinguishing between healthy host tissue and foreign materials (*e*.*g*., bacteria, microbes, antigens, etc.), functioning as a surveillance system to uphold homeostasis and respond to potential threats.^17^ Activation of the complement system on foreign surfaces can occur through three different pathways: the classical, lectin, or alternative pathways.^16^ The C1 complement proteins (*e*.*g*., C1qa, C1qb, and C1qc) are synthesized by macrophages and monocytes, and play an important role in the activation of the classical pathway in collaboration with host antibody-mediated recognition. Activation of the complement cascade by all three pathways leads to the cleavage of a central complement mediator, C3, into fragments C3a and C3b.^18^ These fragments combine with other complement components and form active complexes, such as C5 convertase, that initiates dissociation of C5 into its C5a and C5b fragments. These fragments are strong activators and chemoattractants for neutrophils, macrophages, and other promoters of inflammation.^10,19,20^ Specifically, C5a is a strong activator of myeloid leukocytes via complement receptors C5aR1 and C5aR2.^21^ It has been shown that biomaterial-mediated activation of the complement cascade leads to the release of complement C3a and C5a fragments, while complement proteins such as C1q and C3b are often found in the protein adsorbate to the material itself.^22^

Given the canonical role that the complement system plays in the host foreign body response to implanted biomaterials, we were interested in investigating the role of complement activation, or the lack thereof, in subcutaneous implants of MAP scaffolds compared to NP hydrogel. Recent work by Nicklow et al. demonstrated tissue integration and quiescence within MAP scaffolds over a 12-month timespan in a subcutaneous implantation model, providing evidence of long-term stability and further motivating our investigation.^23^ We used mass cytometry (CyTOF), single cell RNA sequencing (scRNA-seq), and ELISA/multiplex cytokine immunoassays to fill this gap in knowledge of how MAP scaffolds evade the foreign body response on a cellular, proteomic, and transcriptomic level. We hypothesized that biomaterial porosity promotes the inhibition of canonical drivers for FBR, including inflammatory-polarized immune cells, pro-inflammatory cytokines, and complement proteins/receptors.

## RESULTS

### CyTOF demonstrates MAP and NP implants are enriched with distinctly different immune cell populations over time

We wanted to characterize the differences in how MAP scaffold and NP hydrogel implants modulate infiltrating cellular responses in immune and stromal cell populations. To do this, we designed a CyTOF panel of 33 metal isotope-labeled antibodies to identify and quantify shifts in scaffold-infiltrating cell populations (**Table S1**). MAP scaffolds or NP hydrogels were subcutaneously implanted by injection into the dorsal side of Swiss Webster mice (n=6 mice per timepoint per condition). After 3, 5, 7, 14, 21, 35, and 49 days, the implants were removed, enzymatically digested into a single cell suspension, and the cells were stained for CyTOF analysis. To identify cell populations, we used a manual gating strategy on our dataset (**Figure S1**). While there were no changes in total immune cell numbers (**Figure 1A**), changes existed among immune cell populations. Neutrophil recruitment peaked early at day 3 as expected with an increased presence among MAP implants (**Figure 1B**), potentially due to increased pore size being able to retain more neutrophils within the implant following excision. Mast cell numbers remained low with the exception of a peak in mast cell recruitment among NP implants at day 7 (**Figure 1C**). This observation is corroborated by previous work which demonstrated that mast cells contribute to fibrotic encapsulation of implanted biomaterials.^24^

**Figure 1:**
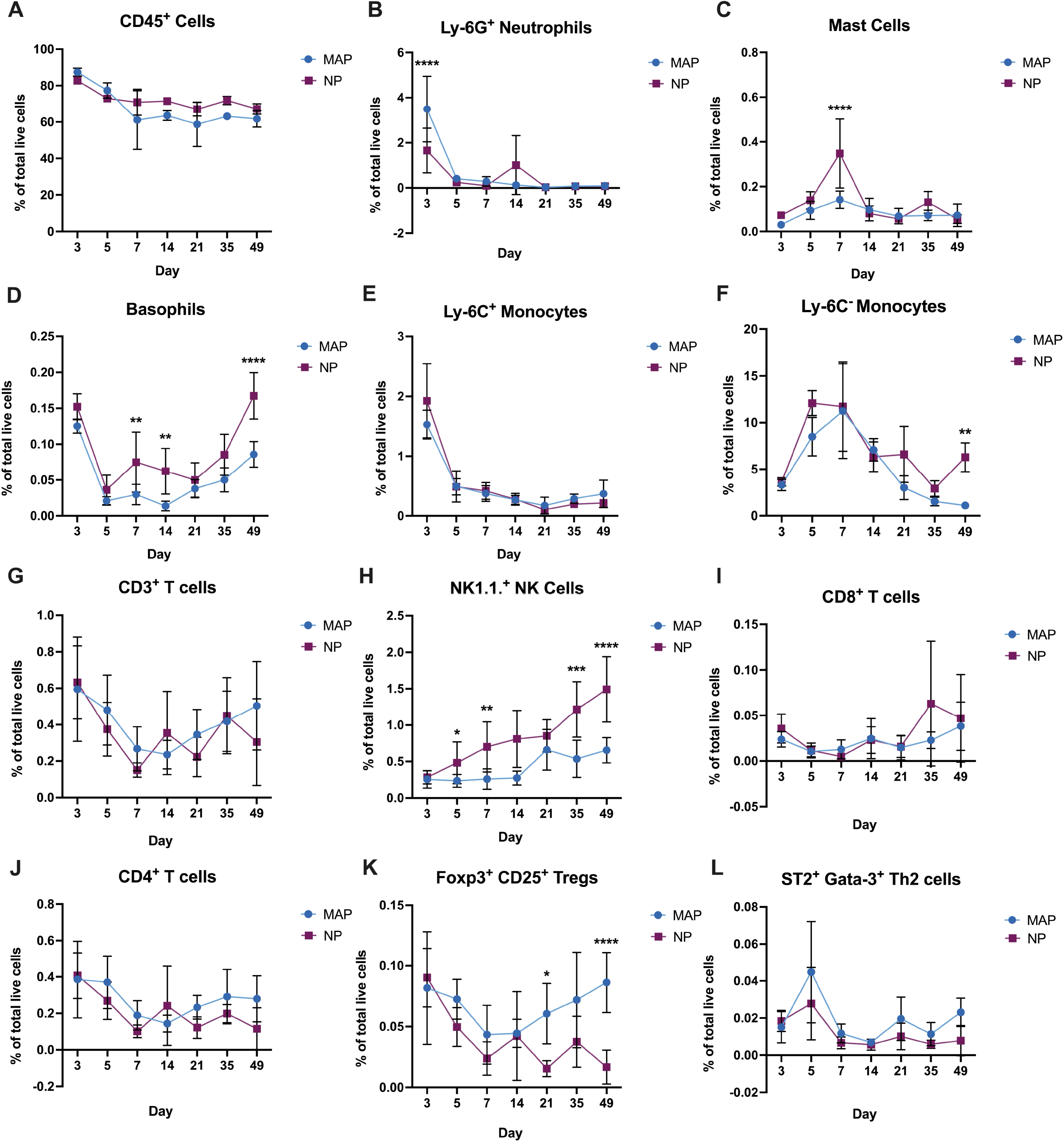
MAP and NP implants are enriched with distinctly different immune cell populations over time. A) Percentage of CD45^+^ live cells within MAP and NP implants over time. B) Percentage of Ly-6G^+^ neutrophils (CD45^+^, CD11b^+^, Ly-6G^+^) within MAP and NP implants over time. C) Percentage of mast cells (CD45^+^, CD11b^-^, CD11c^-^, FceR1a^+^, CD117^+^) within MAP and NP implants over time. D) Percentage of basophils (CD45^+^, CD11b^-^, CD11c^-^ FceR1a^+^, CD117^-^) within MAP and NP implants over time. E) Percentage of Ly-6C^+^ monocytes (CD45^+^, CD11b^+^, F4/80^+^, CD11c^-^ Ly-6C^+^) within MAP and NP implants over time. F) Percentage of Ly-6C^-^ monocytes (CD45^+^, CD11b^+^, F4/80^+^, CD11c^+^, Ly-6C^-^) within MAP and NP implants over time. G) Percentage of CD3^+^ T cells (CD45^+^, CD11b^-^, CD3^+^) within MAP and NP implants over time. H) Percentage of natural killer (NK) cells (CD45^+^, CD11b^-^, CD11c^-^, CD3^+^, NK1.1^+^, FceR1a^-^) within MAP and NP implants over time. I) Percentage of CD8^+^ T cells (CD45^+^, CD11b^-^, CD3^+^, CD4^-^, CD8^+^) within MAP and NP implants over time. J) Percentage of CD4^+^ T cells (CD45^+^, CD11b^-^, CD3^+^, CD4^+^, CD8^-^) within MAP and NP implants over time. K) Percentage of regulatory T (Treg) cells (CD45^+^, CD11b^-^, CD3^+^, CD4^+^, Foxp3^+^, CD25^+^) within MAP and NP implants over time. L) Percentage of T helper 2 (Th2) cells (CD45^+^, CD11b^-^, CD3^+^, CD4^+^, ST2^+^, Gata-3^+^) within MAP and NP implants over time. Statistical analysis: An ordinary two-way ANOVA with Sidak’s multiple comparisons test was used to compare differences between groups at each timepoint for all datasets. Significance symbols: * p < 0.05, ** p < 0.01, *** p < 0.001, **** p < 0.0001. Error bars: mean ± SD per group. Outlier analysis: Outliers from any of the datasets were removed using the Robust Regression and Outlier Removal (ROUT) method (Q = 1%) in GraphPad Prism.

Cell populations were dynamic for both the NP and MAP groups, notably there was a progressive and significant increase in basophils (**Figure 1D**) within the NP implants that did not clear from the implant site over time. Basophils are canonically known for their role in the allergy response^25^, but in the context of implanted biomaterials, activated basophils have been shown to upregulate IL-4 and IL-13 and their sustained activation and presence has been implicated in fibrotic encapsulation, which aligns with our findings.^26^ While there were little to no differences between MAP and NP implants in monocytes (**Figures 1E-F**) and T cells (**Figure 1G**), there was a noteworthy progressive increase in natural killer (NK) cells throughout the course of the study starting from the day of implantation that was significantly higher in NP implants (**Figure 1H**). NK cells secrete IFN-γ, TNF-α, and IL-1β which enhance macrophage fusion on biomaterials and may impede tissue integration.^27,28^ Among T cell populations, there were differences between MAP and NP implants in CD8+ cytotoxic T cells (**Figure 1I**), CD4+ helper T cells (**Figure 1J**), and Th2 T cells (**Figure 1L**). Interestingly, while CD4+ helper T cells remained constant within the MAP scaffold implants, there was a significant progressive increase in Foxp3^+^ regulatory T cells (Tregs) over time (**Figure 1K**), indicating a decrease in other helper T cell populations. Tregs are known to have anti-inflammatory, tolerogenic effects on the host immune response to biomaterials and promote tissue repair and regeneration.^29^ Altogether, these data demonstrate the key differences in immune cell populations among fibrotic NP implants and non-fibrotic MAP implants.

We next sought to determine differences among macrophage and dendritic cells between NP and MAP implants. There were interesting fluctuations in macrophage polarization between the hydrogel implants over time. Approximately 50-70% of all cells within these implants were F4/80^+^ macrophages throughout the study (**Figure 2A**) while dendritic cells comprised 0-2% of all cells (**Figure 2B**). Nearly half of the macrophages were CD86^+^ CD206^-^ “M1” polarized within the first 3-5 days post-implant (DPI), which was likely due to the initial inflammation from the hydrogel implantation (**Figure 2C**).^30^ MAP implants displayed an elevated CD86+ M1 macrophage population at days 3 and 5 after implantation while NP implants demonstrated a brief elevated presence of M1 macrophages out at day 21, demonstrating the ability of MAP implants to resolve acute M1 macrophage-mediated inflammation. There was also a slight increase in CD86^-^ CD206^+^ “M2” macrophages over time, and the NP implants contained significantly higher numbers of M2 macrophages compared to the MAP scaffolds (**Figure 2D**). While M2 macrophages are known as the more anti-inflammatory / pro-regenerative phenotype, certain M2 subtypes are potent modulators of IL-4 and IL-13 cytokine secretion, which can lead to FBGC formation and the formation of a foreign body capsule around the implanted biomaterial.^31^ The nearly 2-fold increase in the percentage of M2 macrophages within the NP implants and sustained higher levels of M2 macrophages with NP implants indicates that prolonged presence of M2 macrophages is associated with the FBR seen against NP implants. There is also an increase of CD86^+^ CD206^+^ “M++” macrophages (**Figure 2E**) and a simultaneous decrease of CD86^-^ CD206^-^ “M0” macrophages (**Figure 2F**) over time. This indicates that there are macrophages undergoing activation that are not falling within the M1 and M2 macrophage activation states. Given evidence that macrophage activation is a gradient^32^, it is likely that these M0 macrophages may adopt an activation state that is not captured by the M1 and M2 macrophage paradigm yet is present within the M++ macrophage population.

**Figure 2:**
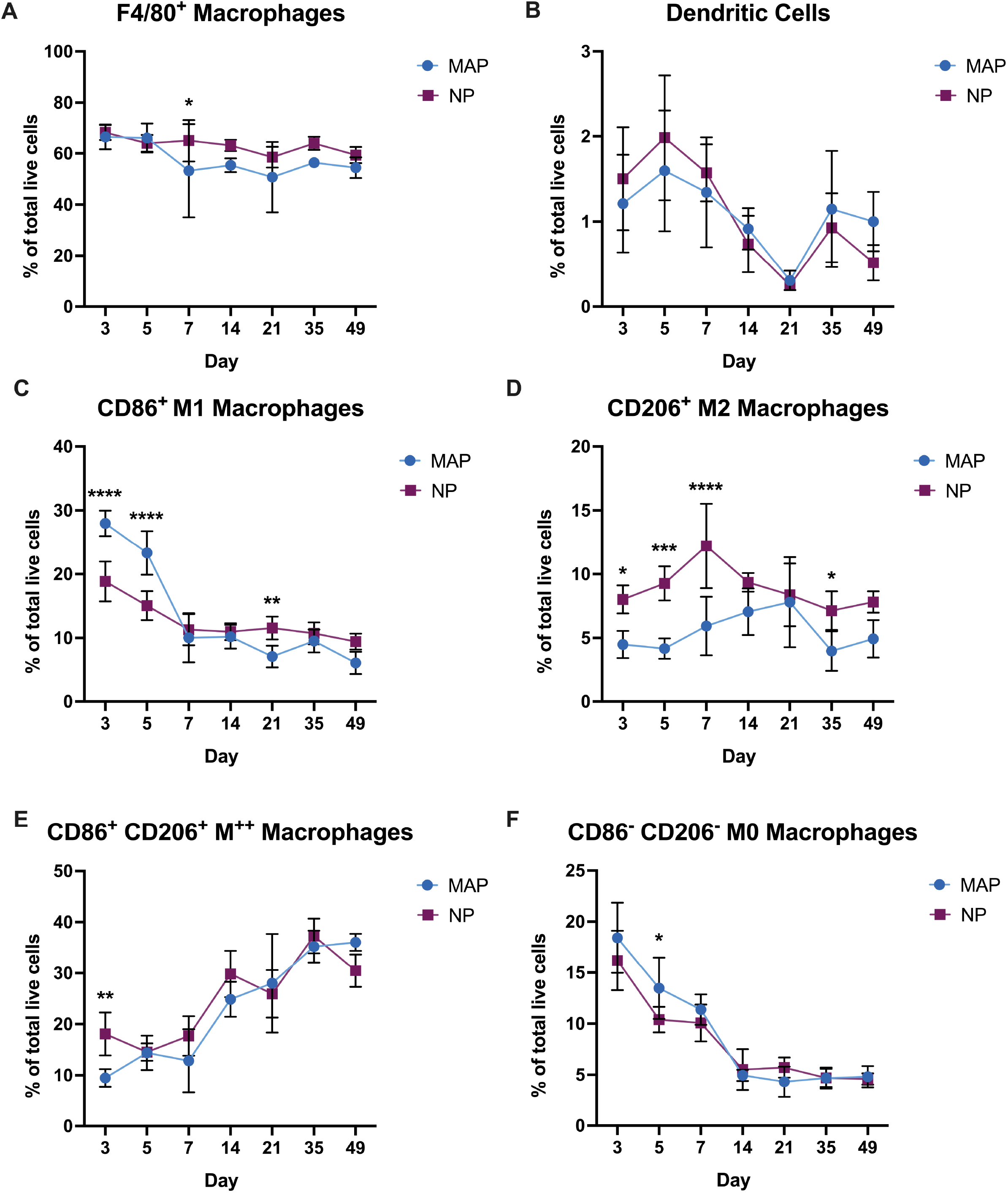
Macrophage populations in MAP and NP implants dynamically fluctuate over time. A) Percentage of macrophages (CD45^+^, CD11b^+^, F4/80^+^) within MAP and NP implants over time. B) Percentage of dendritic cells (CD45^+^, CD11c^+^, F4/80^+^, MHCII^+^, CD54^+^) within MAP and NP implants over time. C) Percentage of M1 macrophages (CD45^+^, CD11b^+^, F4/80^+^, CD86^+^, CD206^-^) within MAP and NP implants over time. D) Percentage of M2 macrophages (CD45^+^, CD11b^+^, F4/80^+^, CD86^-^, CD206^+^) within MAP and NP implants over time. E) Percentage of M^++^ macrophages (CD45^+^, CD11b^+^, F4/80^+^, CD86^+^, CD206^+^) within MAP and NP implants over time. F) Percentage of M0 macrophages (CD45^+^, CD11b^+^, F4/80^+^, CD86^-^, CD206^-^) within MAP and NP implants over time. Statistical analysis: An ordinary two-way ANOVA with Sidak’s multiple comparisons test was used to compare differences between groups at each timepoint for all datasets. Significance symbols: * p < 0.05, ** p < 0.01, *** p < 0.001, **** p < 0.0001. Error bars: mean ± SD per group. Outlier analysis: Outliers from any of the datasets were removed using the Robust Regression and Outlier Removal (ROUT) method (Q = 1%) in GraphPad Prism.

We next sought to determine shifts in cell populations outside the immune compartment between MAP and NP hydrogel implants. Among CD45^-^ cells, MAP and NP implants demonstrated a similar increase over time (**Figure 3A**). Additionally, both MAP and NP hydrogel implants exhibited a similar increase in abundance of total CD140a^+^ CD49b^+^ fibroblasts over time with an increased presence among NP implants only at 3 DPI (**Figure 3B**). Looking further within the fibroblast compartment, there was a significant increase in αSMA^+^ Collagen I^+^ myofibroblasts in the MAP scaffolds 5 and 7 DPI (**Figure 3C**), which are cells that play an important role in tissue repair. While accumulation of these cells can result in collagen deposition that may lead to fibrosis, the myofibroblasts were quickly cleared from the MAP implant site. There was an increase in CD326+ epithelial cells in MAP implants at 7 DPI (**Figure 3D**), potentially indicating the onset of a re-epithelialization response among the regenerative MAP implants. Additionally, there was a significant increase in CD31^+^ endothelial cells in the MAP scaffolds at 14 DPI and onward compared to NP hydrogel implants (**Figure 3E**), once again indicating that the ingrowth of new vasculature is dependent on sufficient material porosity.

**Figure 3:**
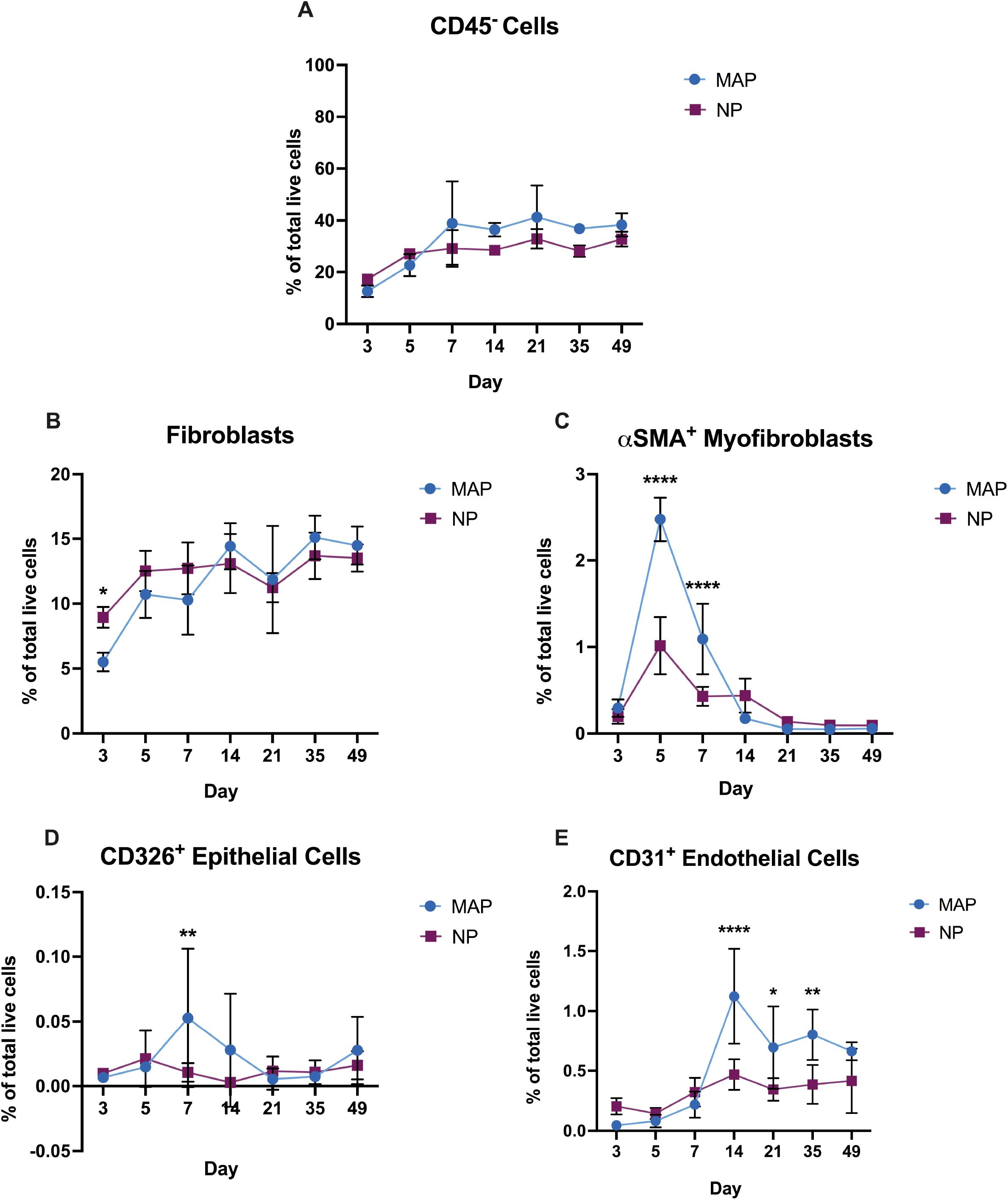
Enrichment of stromal cell populations within MAP implants over time suggest improved tissue implant integration compared to NP implants. A) Percentage of CD45^-^ live cells within MAP and NP implants over time. B) Percentage of fibroblasts (CD45^-^, CD11b^-^, CD140a^+^, CD49b^+^) within MAP and NP implants over time. C) Percentage of myofibroblasts (CD45^-^, CD11b^-^, CD140a^+^, CD49b^+^, col1a1^+^, αSMA^+^) within MAP and NP implants over time. D) Percentage of epithelial cells (CD45^-^, CD140a^-^, CD326^+^) within MAP and NP implants over time. E) Percentage of endothelial cells (CD45^-^, CD140a^-^, CD31^+^) within MAP and NP implants over time. Statistical analysis: An ordinary two-way ANOVA with Sidak’s multiple comparisons test was used to compare differences between groups at each timepoint for all datasets. Significance symbols: * p < 0.05, ** p < 0.01, *** p < 0.001, **** p < 0.0001. Error bars: mean ± SD per group. Outlier analysis: Outliers from any of the datasets were removed using the Robust Regression and Outlier Removal (ROUT) method (Q = 1%) in GraphPad Prism.

### scRNA sequencing elucidates differential expression in complement-associated proteins between MAP and NP implants

While our CyTOF analysis identified noteworthy differences in cell infiltration within the MAP and NP hydrogel implants over a 49-day period, we wanted a more in-depth analysis of transcriptome-level differences of single cells isolated from MAP and NP hydrogel implants after 7 and 21 days post-implant (DPI).

After 7 and 21 days, subcutaneous MAP and NP implants were isolated, digested into a single-cell suspension, and cDNA libraries were constructed using a 10X Genomics Chromium X platform followed by single cell RNA-sequencing (scRNA-seq). We performed downstream analysis of the sequencing data using Seurat v5^33,34^. We created Uniform Manifold Approximate Projection (UMAP) plots for each implant type at each timepoint (**Figure 4A-B**) and annotated cell types according to validated literature marker genes (**Figure 4C-D**). The relative distribution of cell subpopulations matched those reported by other publications characterizing cellular responses to subcutaneous biomaterial implants^35–37^. One pathway long attributed to inflammatory responses and that informs the FBR to implanted biomaterials is the complement pathway^38^, yet it remains unknown whether the regenerative capacity of MAP implants is due to modulation of the complement pathway. Therefore, we used our scRNA-seq dataset to investigate the role complement activation plays in the FBR to our biomaterial implants. Among 7 DPI macrophages (*Ptprc+Adgre1+Cd68+*), C1qb and C1qc were significantly upregulated in NP gels, whereas C5ar1 was upregulated in MAP gels (**Figure 4E**). Conversely, in 21 DPI samples, C1qa and C1qb were upregulated in MAP, while the downstream complement receptors were upregulated in NP. It is noteworthy that components of the complement protein C1 were upregulated in 7 DPI NP samples, as it is a main player in the activation of the classical complement pathway. This pathway initiates a response against foreign materials and can lead to subsequent activation of C3a and C5a, which are the main pro-inflammatory anaphylatoxins of the complement system.^16,39^ Moreover, while there were increased C1 components among MAP implants 21 DPI, previous work has demonstrated the ability of C1q complement proteins of promoting a pro-resolving macrophage phenotype in other contexts.^40^ Additionally, macrophages found within NP implants displayed a higher upregulation of complement receptors, potentially indicating a heightened responsiveness to complement activation not present in MAP gels. Given this interesting result, we wanted to investigate this further and determine which cell subpopulation(s) were responsible for expression of complement components and complement receptors.

**Figure 4.**
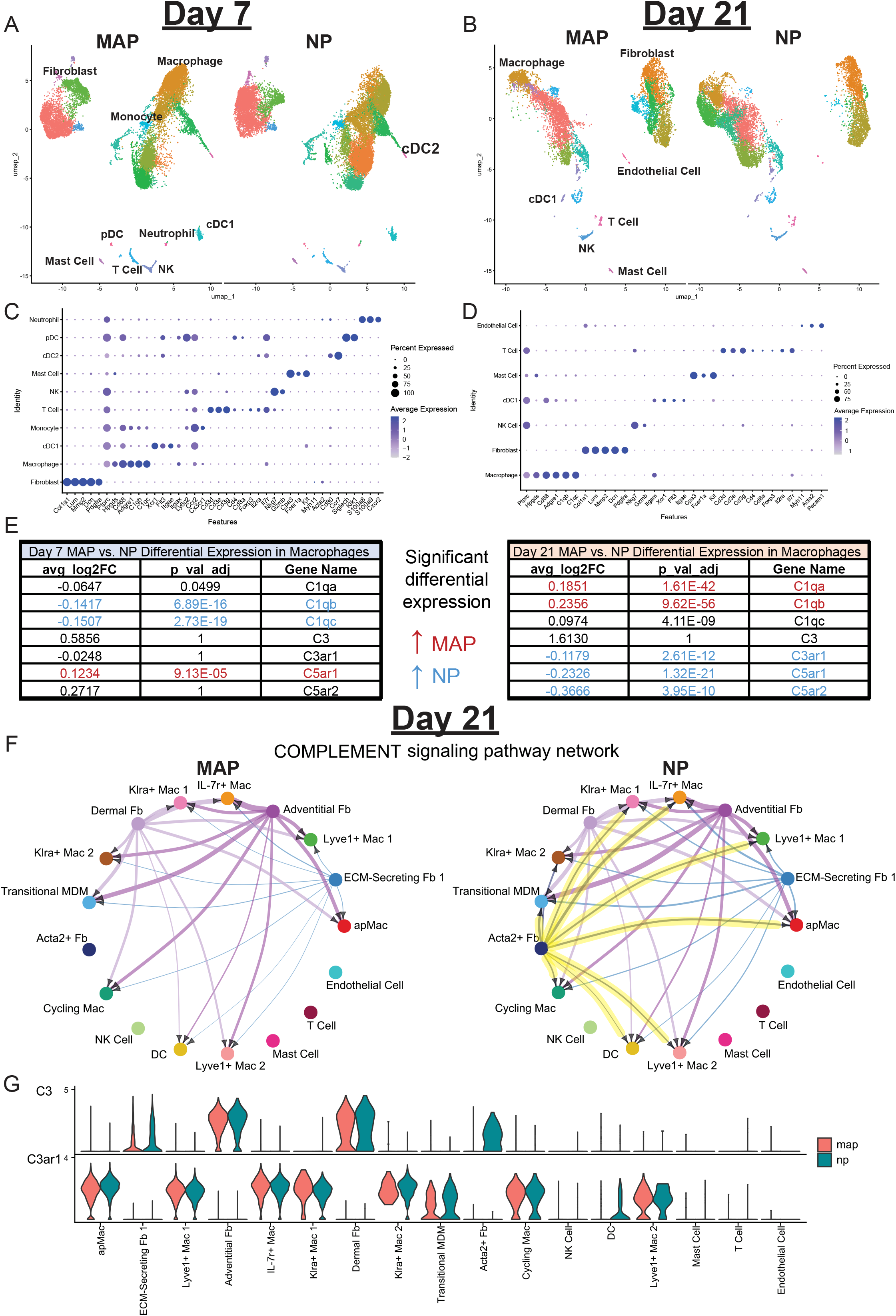
Differential gene expression between MAP and NP implants. A) Uniform Manifold Approximate Projection (UMAP) plot displaying cells clustered by cell type for 7 and B) 21 days post-implant (DPI). Each plot is separated into MAP- and NP-exclusive cells. (C and D) Dot plot displaying canonical cell type markers for each cell type for 7 (C) and 21 (D) DPI. E) Table showing differential expression of complement pathway-associated genes in 7 (left) and 21 (right) DPI with associated adjusted p values (p_val_adj) and average log_2_ fold changes (avg_log2FC). (F) Cell-cell communication circle plots of the complement pathway among cells from MAP (left) and NP (right) gels 21 DPI. Arrowheads indicate which cell subpopulation is the receiver for each interaction. (G) Violin plot depicting expression of the C3 and C3ar1 genes across cell subpopulations in MAP and NP samples 21 days post-implant. ap = antigen-presenting; Mac = macrophage; Fb = fibroblast; MDM = monocyte-derived macrophage; NK = natural killer; DC = dendritic cell; pDC = plasmacytoid dendritic cell; cDC = classical dendritic cell.

We conducted further analysis using the CellChat^41^ package (v1.6) to identify the cell subpopulations engaging in differential complement signaling 21 DPI. We determined that Acta2+ fibroblasts displayed elevated expression of complement components that were predicted to engage with different macrophage subpopulations based on elevated complement receptor expression, all exclusively in NP implants, indicating a potential myofibroblast-macrophage axis (**Figure 4F**). A closer look at C3-C3ar1 signaling highlighted an increase in C3 expression across all fibroblasts in NP gels (**Figure 4G**). We then focused on complement signaling between individual fibroblast and immune subpopulations, where C3-C3ar1 signaling was upregulated almost exclusively in NP cell subpopulations (**Figure S2**). This C3-C3ar1 signaling axis aligns with the upregulation of C5 receptors (*C5ar1* and *C5ar2*) identified among macrophages based on previous work demonstrating that C3 signaling leads to inflammatory signaling that promotes elevated expression of C5 receptors in different immune cell populations, including macrophages.^42,43^ Overall, our single-cell analysis revealed the complement pathway as a potential mechanism for the different healing outcomes associated with MAP and NP implants.

Intrigued by these differences, we decided to run an additional experiment where we implanted mice (Swiss Webster strain) with subcutaneous injections of MAP scaffold or NP hydrogel implants (n=3 per group) and removed the implants after 21 days. These implants were then enzymatically digested and the cells were lysed with a lysis buffer to isolate any proteins and/or cytokines from the implant. This cell lysate was then analyzed for C5a and C3a complement proteins, which are commonly activated in the FBR to biomaterials^20,21^, via ELISA (**Figure 5**). There was a significant reduction in C5a protein in the MAP scaffolds compared to the hydrogel implants, which is one of the most potent mediators of the complement pathway and the recruitment of inflammatory immune cells (**Figure 5A)**.^19,22^ There was no difference between amount of C3a protein in the MAP and NP implants (**Figure 5B)**. While both proteins play an important role in the activation of the complement cascade, C5a has been shown to be a 50-100 times more potent chemoattractant than C3a.^44^ This data demonstrates that MAP lowers C5a complement activation.

**Figure 5.**
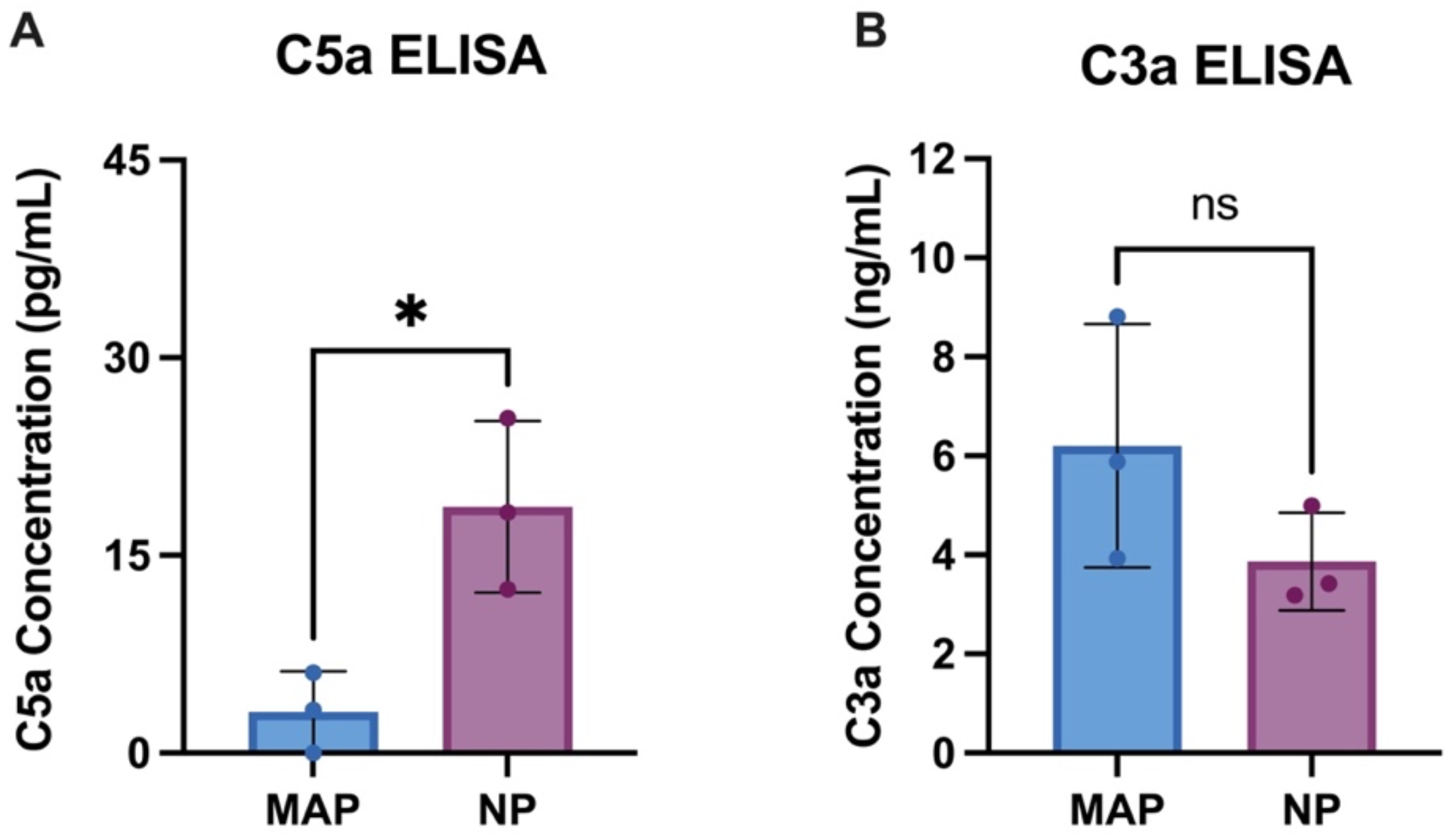
Lower levels of C5a complement protein, but not C3a, are detected in MAP implants compared to NP implants. A) Concentration (pg/mL) of C5a complement protein in MAP and NP implant cell lysates at 21 days post-implant. B) Concentration (ng/mL) of C3a complement protein in MAP and NP implant cell lysates at 21 days post-implant. Statistical analysis: Unpaired parametric t-test was used between MAP (n=3) and NP groups (n=3). Significance symbols: * p < 0.05, ** p < 0.01, *** p < 0.001, **** p < 0.0001. Error bars: mean ± SD per group.

### Inflammatory response and fibrotic encapsulation of NP implants is C5 complement-dependent

To further assess whether the inflammatory and fibrotic nature of NP scaffolds is dependent on activation of C5a, both MAP and NP hydrogels were implanted into C5 deficient (C5 KO; *Hc*^*-/-*^) mice (B10.D2-Hc^0^ H2^d^ H2-T18^c^/oSnJ strain) and wildtype (WT) mice (C57BL/6 strain). After 21 days, the implants were removed and underwent both histological and soluble factor analysis. To capture soluble factors in these implants, hydrogel implants were harvested and digested, and the cell lysate was assessed for inflammatory cytokines via a 32-plex cytokine multiplex assay (Luminex). Within the wildtype mice, there was a significant increase in proinflammatory cytokines and chemokines IL-1β, IL-6, KC, MCP-1, MIP-1β, and TNFα in the NP hydrogel implants compared to the MAP scaffold implants (**Figure 6A**). IL-6, TNFα, IL-1β are canonical inflammatory cytokines that are released by immune cells and, in conjunction with C3a/C5a, can contribute to the exacerbation of the foreign body response to biomaterials.^45^ MCP-1 and MIP-1β are inflammatory chemokines that signal for macrophage, monocyte, and NK cell trafficking.^46,47^ Additionally, there has been reported evidence that increased MCP-1/CCL2 participates in macrophage fusion and foreign body giant cell formation.^31,48^ IL-5 and IL-9, which are classified as Th2 cytokines that are typically anti-inflammatory in nature, were significantly more enriched in MAP scaffolds in WT mice.^49^

**Figure 6.**
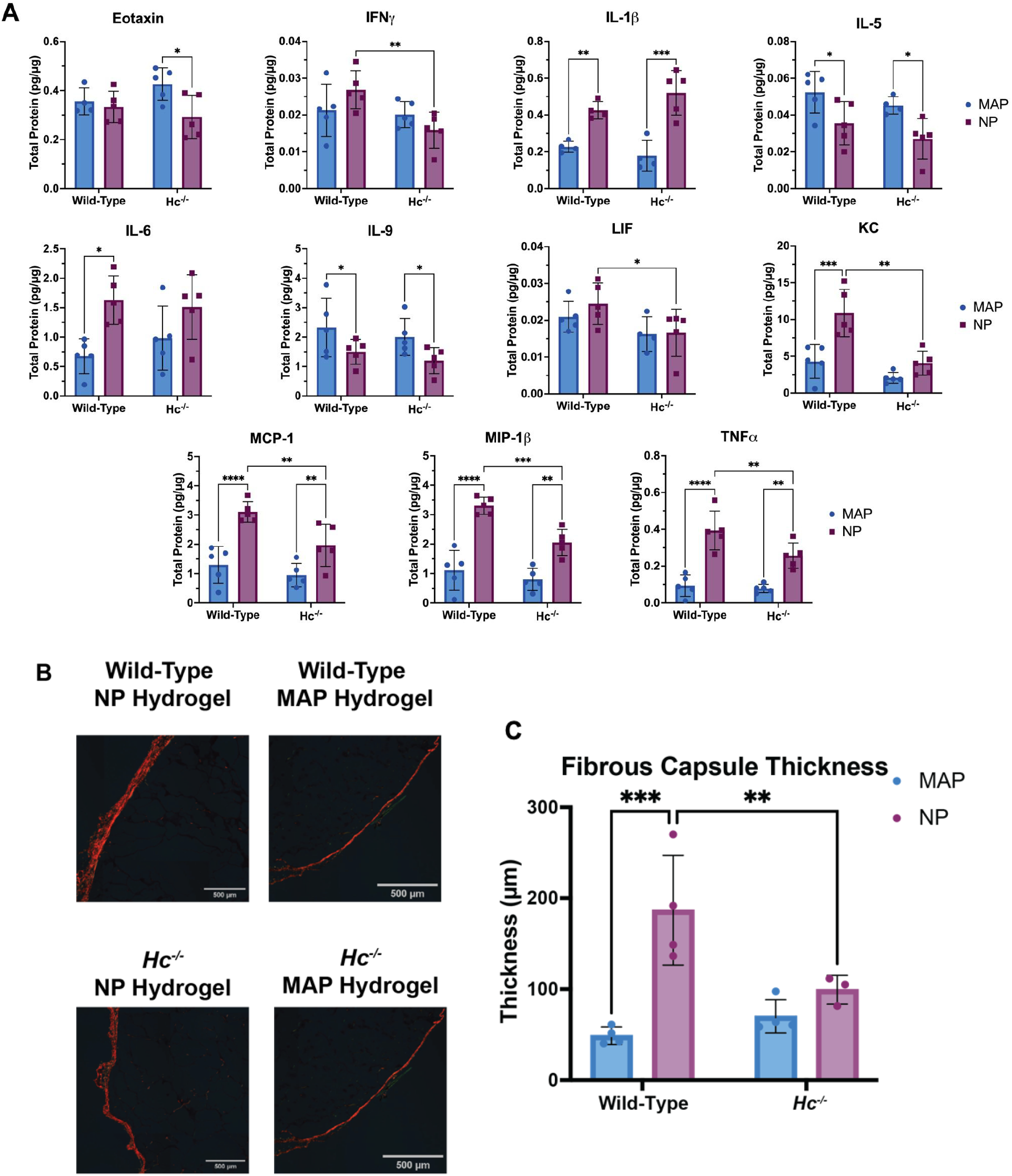
Inflammatory response and fibrotic encapsulation of NP implants is C5 complement-dependent. A) A cytokine multiplex assay was used to detect cytokines within MAP and NP implants harvested from wild-type and C5 deficient (HC^-/-^) mice. Statistical analysis: An ordinary two-way ANOVA with Sidak’s multiple comparisons test was used to compare differences between groups at each timepoint for all datasets. B) Picrosirius red staining of hydrogel implants imaged under polarized light. C) Thickness of fibrous capsule measurements taken across all implant samples. Significance symbols: * p < 0.05, ** p < 0.01, *** p < 0.001, **** p < 0.0001. Error bars: mean ± SD per group.

Interestingly, in the C5 KO/*Hc*^*-/*-^ mice there was a significant decrease in inflammatory cytokines IFNγ, TNFα, and LIF along with significant decreases in the chemokines KC (i.e. CXCL1), MCP-1, and MIP-1β, within the NP implants compared to the wildtype mice (**Figure 6A)**. These data align with the known role of C5a as a potent chemoattractant that promotes macrophage recruitment to the site of pathogens or foreign objects. Furthermore, histological analysis using picrosirius red staining was conducted under polarized light to identify collagenous deposition, including the identification of fibrotic capsule formation around implanted hydrogels (**Figure 6B**). This histology analysis determined that MAP implants elicited a significantly thinner collagen capsule than NP implants in wild-type mice, which matches previously shown work.^13^ However, the capsule of NP implants in C5 KO/*Hc*^*-/*-^ mice is significantly reduced compared to wild-type and not statistically different than MAP in either genotype (**Figure 6C**). Given that C5a and C5b are key players in orchestrating the foreign body response to biomaterial implants, these data confirm that C5 plays a large role in the inflammatory and fibrotic response to NP implants that is dampened by MAP implants. Therefore, MAP scaffolds elicit a lower level of C5a activation which could contribute to its ability to evade the foreign body response. The C5 KO cytokine and picrosirius red data, in combination with the previous results in this manuscript, provides rigorous support for a simple and temporally robust roadmap to biomaterial avoidance of complement activation and, subsequently, on the foreign body response to long-term implants enabled by the MAP technology.

## DISCUSSION

In this manuscript, we used CyTOF, scRNA-sequencing, and ELISA/multiplex cytokine immunoassays to investigate the cellular, proteomic, and transcriptomic differences in the host immune response to subcutaneous implants of MAP scaffold and nanoporous (NP) hydrogels.

The CyTOF data identified infiltrating immune and stromal cell populations within the implants at 3, 5, 7, 14, 21, 35, and 49 days post-implant (DPI). Notably, there was a significant increase in CD31^+^ endothelial cells in the MAP scaffolds 14 DPI and onward compared to NP hydrogel implants, which is consistent with published findings that the porosity of MAP scaffold is conducive for angiogenesis.^14^ Additionally, there was a significant progressive increase in Foxp3+ Tregs over time in the MAP scaffolds. These cells are known to have anti-inflammatory, tolerogenic effects on the host immune response to biomaterials and promote tissue repair and regeneration.^29^ Within the NP implants, there was a progressive increase in basophils and NK cells that did not clear from the implant site over time. Basophils are canonically known for their role in allergic response^25^, while NK cells secrete IFN-γ, TNF-α,and IL-1β which enhances macrophage fusion on biomaterials and may impede tissue integration.^26,27^ There was also a significant increase in CD86-CD206+ “M2” macrophages in NP implants at early timepoints (3-7 days) compared to the MAP scaffolds. Certain M2 sub-phenotypes are potent modulators of IL-4 and IL-13 cytokine secretion, which can lead to foreign body giant cell (FBGC) formation and fibrosis of the implanted biomaterial.^31^

The scRNA-seq data elucidated interesting differences in complement pathway activation between MAP and NP implants at 7 and 21 DPI. The foreign body response (FBR) is canonically exacerbated by the activation of the complement pathway, which is a group of sequentially reacting proteins that mediate various biological processes that play an important role in host immune response.^16^ It has been shown that biomaterial activation of the complement cascade leads to the release complement C3a and C5a fragments, while complement proteins such as C1q and C3b are often found in the protein adsorbate.^22^ Differential gene expression between MAP and NP implants at 7 and 21 DPI showed that there was a significant downregulation of genes encoding for C1 proteins and complement C5a receptor in certain macrophage subpopulations. ELISA data further supported this result, as there was a significant decrease in C5a protein content in MAP scaffold implants compared to NP implants. Since C5a is one of the most potent chemoattractants in the complement cascade^44^, it may be a driving force in the FBR to nanoporous implants. This was investigated further by utilizing a C5 KO/*Hc*^*-/*-^ mouse model.

A Luminex multiplex cytokine assay was run on cell lysate from MAP and NP implants at 21 DPI in C5 KO/*Hc*^*-/*-^ mice and WT mice. Relative to the MAP scaffold implants, there was a significant increase in proinflammatory cytokines IL-1β, IL-6, KC, MCP-1, MIP-1β, and TNFα in the NP hydrogel implants in WT mice. All of these cytokines play a role in immune cell trafficking and foreign body giant cell formation. In the C5 KO/*Hc*^*-/*-^ mice, a decrease in cytokine concentration was observed only for the NP implants for IFNγ, KC, MCP-1, MIP-1β, and TNFα, which are key players in orchestrating the FBR. There was no difference observed in the cytokines for MAP implants between the C5 KO/*Hc*^*-/*-^ and WT mice, indicating that the inflammatory response to MAP (or lack thereof) is not dependent on C5 activation.

One limitation of this work is that a portion of our data is at the transcript level and regulation of the complement pathway is post-translational. While transcript level regulation does not always equate to protein level differences, our ELISA data demonstrates complement activation at the protein level and our *in vivo* work demonstrates the functional relevance of complement in the fibrotic encapsulation of implanted biomaterials. Further work is needed in to determine how transcript level complement activation is regulated.

All these data collectively provide supporting mechanistic evidence to support our hypothesis that the cell-scale porosity of MAP scaffolds acts by reducing canonical drivers for FBR, including inflammatory-polarized immune cells and pro-inflammatory cytokines. Further and mechanistically, this data provides protein and RNA-level evidence that this avoidance is, in part, due to diminished activation of C5 complement pathway. Altogether these results show that MAP scaffold porosity down regulates the complement-fibroblast-macrophage signaling loop, revealing a key mechanism through which MAP scaffolds mitigate the foreign body response and provide the capability of previously reported prolonged tissue integration.^23,50^

## METHODS

### MAP microgel synthesis

The 5.0 wt% (w/v) microgel precursor solution consisted of 73.2 mg mL^-1^ 4-arm PEG-maleimide with 0.004 M excess of unreacted maleimides (10 kDa, Nippon Oil Foundry, Japan), 12.2 mg mL^-1^ MMP-2 degradable crosslinker (Ac-GCGPQGIAGQDGCG-NH2, Watson Bio), 0.75 mg mL^-1^ RGD (Ac-RGDSPGGC-NH2, Watson Bio), 5 µM of biotin-maleimide, and 8.02 mg mL^-1^ MethMal annealing macromer. Microgels were synthesized using a high-throughput microfluidics technique as previously described.^51^ The microgel precursor solution was run at 3 mL hr^-1^ and the surfactant solution (2% Pico-Surf in NOVEC 7500) was run at 6 mL hr^-1^ through a PDMS microfluidic device (parallelized step-emulsification) with a step size of 11.3 µm. After gelation, the microgels were removed from the oil phase by washes with oil, 1X PBS (pH 3.0), and hexanes as previously described. Microgels were then sterilized with three 70% IPA washes followed by three washes with sterile 1X PBS (pH 3.0). A PBS buffer with an acidic pH was used so that the unreacted maleimides were not susceptible to hydrolysis. To inactivate the excess maleimides, the microgels were incubated with a strong base, triethylamine (20 µL per 1 mL of gel) overnight (inactivated MAP), and then washed with 1X PBS (pH 7.4) to remove the triethylamine. Nanoporous gels were made with a chemically identical 5.0 wt% formulation, but the PEG-MAL, RGD, and MethMal were dissolved in pH = 4.3 10X PBS and the MMP-2 crosslinker was dissolved in pH = 7.4 1X PBS. The PEG-MAL solution and crosslinker solution was combined at a 1:1 volume and gelation time was confirmed by rheology to be approximately 7 minutes.

### Subcutaneous injections of MAP scaffolds into Swiss Webster mice

All animal work was approved by the University of Virginia Animal Care and Use Committee (Protocol #4165. For these experiments, 8-week-old, female Swiss Webster mice (Charles River Laboratories) were used. The mice were anesthetized with isoflurane administered with oxygen before any procedure. Prior to the injections, the dorsal side of each mouse was depilated by shaving and applying hair removal cream (Nair) then cleaning the skin with an alcohol wipe. The MAP microgels were incubated 1:1 (v/v) with a sterile 40 µM Eosin Y solution for at least 15 min, spun down at 18,000 x g for 5 min to remove excess Eosin Y, and then loaded into a sterile 1 mL syringe with a positive displacement pipette. The microgels were injected into the subcutaneous space under the skin (50 µL volume, n=4 implants per mouse) with a 23-guage needle and each implant was irradiated with 505 nm LED light (ThorLabs, 8.7 mW cm^-2^ intensity) for 1 min to anneal the scaffolds under the skin. The nanoporous hydrogel precursor components were mixed at a 1:1 ratio approximately 5 minutes prior to injection and then injected into the subcutaneous space under the skin (50 µL volume, n=4 implants per mouse) approximately 1 minute before gelation began. Since the gelation is pH dependent, the nanoporous (NP) hydrogels fully crosslink underneath the skin which has a relatively neutral pH. Each mouse received only one type of implant, MAP or NP, (n=4 implants per mouse), and there were 6 mice per group per timepoint for the CyTOF experiment (3, 5, 7, 14, 21, 35, and 49 days), 3 mice per group per timepoint for the scRNA-seq experiment (7 and 21 days), and 3 mice per group for the C5a/C3a ELISA experiment (21 day timepoint).

### Subcutaneous injections of MAP scaffolds into C5 deficient (B10.D2) mice

All animal work was approved by the University of Virginia Animal Care and Use Committee (Protocol #4165. For this experiment, 12-week-old, female C57BL/6 wildtype (WT) mice (Jackson Laboratories) and 12-week-old, female B10.D2 (B10.D2-Hc^0^ H2^d^ H2-T18^c^/oSnJ strain) mice (Jackson Laboratories), which are serum C5 deficient (C5 KO/*Hc*^*-/*-^), were used. The mice were anesthetized with isoflurane administered with oxygen before any procedure. Prior to the injections, the dorsal side of each mouse was depilated by shaving and applying hair removal cream (Nair) then cleaning the skin with an alcohol wipe. The MAP microgels were incubated 1:1 (v/v) with a sterile 40 µM Eosin Y solution for at least 15 min, spun down at 18,000 x g for 5 min to remove excess Eosin Y, and then loaded into a sterile 1 mL syringe with a positive displacement pipette. The microgels were injected into the subcutaneous space under the skin (50 µL volume, n=4 implants per mouse) with a 23-gauge needle and each implant was irradiated with 505 nm LED light (ThorLabs, 8.7 mW cm^-2^ intensity) for 1 min to anneal the scaffolds under the skin. The nanoporous hydrogel precursor components were mixed at a 1:1 ratio approximately 5 minutes prior to injection and then injected into the subcutaneous space under the skin (50 µL volume, n=4 implants per mouse) approximately 1 minute before gelation began. Since the gelation is pH dependent, the nanoporous (NP) hydrogels fully crosslink underneath the skin which has a relatively neutral pH. Each mouse (WT or C5 KO) received only one type of implant, MAP or NP, (n=4 implants per mouse), and there were five mice per group, for a total of 20 mice. The implants were harvested after 21 days post-implantation.

### Cell isolation from implants

Surgical scissors and forceps were used to remove the implants from the subcutaneous space or the entire fat pad containing the implant. Implants were placed into a tube containing 1 mL collection buffer (RPMI media containing 10 % FBS) and stored on ice. Implants were then added to 1 mL of digestion buffer containing 100 µg mL^-1^ Liberase TM (Roche) and 10 U mL^-1^ DNAse (New England BioLabs) diluted in serum-free RPMI media. Implants were incubated in digestion buffer for 30 min at 37 °C on a tube rotator. After complete degradation of the hydrogel implants, the solution was filtered with 70 µm cell strainer to isolate the cells.

### Cell staining for CyTOF

3×10^6^ cells per mouse sample were added to 1.2 mL low-binding bullet tubes, spun down at 300 x g, and resuspended in cell staining buffer (Standard BioTools) with 3% mouse Fc block (Biolegend). After blocking for 30 min, the cells were spun down, washed, and resuspended in 100 µL of the surface antibody cocktail. After 30 min staining period, the cells were spun down, washed, and stained with cisplatin (live/dead stain). Cells were then fixed and permeabilized using a Foxp3 staining buffer kit (Invitrogen) for 30 minutes. Cells were then resuspended in 100 µL of an intracellular antibody cocktail. The antibodies used in this CyTOF panel are outlined in **Table S1**. Antibodies were purchased from Standard BioTools or conjugated to the isotopes in-house with a Standard BioTools Maxpar Labeling Kit. After 30 min staining period, the cells were spun down, washed, and barcoded with a palladium barcode kit (Standard BioTools) according to manufacturer protocol.^52^ Cells were then intercalated with 125 µM of Cell-ID Intercalator (Standard BioTools). All samples from each timepoint, now assigned a unique barcode, were combined into one tube and frozen in freezing media (90% FBS with 10% DMSO). Before running the samples, each tube was thawed, washed with acquisition solution (Standard BioTools), and then run on a CyTOF 2-Helios cytometer by the University of Virginia Flow Core Facility.

### Debarcoding and analyzing CyTOF data

The data was normalized with beads according to Standard BioTools protocols, then debarcoded using a previously published code. The debarcoded files were imported into OMIQ software (Dotmatics, USA). For the high dimensional data reduction, 2.5×10^6^ cells were subsampled from the dataset and ran through UMAP and FSOM clustering algorithms using OMIQ software. For the manual gating strategy, gates were hand-drawn on cell populations as outlined in **Figure S1**. The data from the gates were exported to GraphPad Prism to construct the graphs and conduct statistical analyses.

### scRNA-seq analysis

At each timepoint (7 and 21 days post-implant), the hydrogel implants were removed and enzymatically digested into a single cell suspension as described above. Cells were resuspended in 0.04% UltraPure BSA in PBS (ThermoFisher). Single cells were barcoded and then libraries were prepared using the 10X Genomics Chromium Single Cell 3’ Reagents Kit (v3). cDNA libraries were amplified and then quantified using the Agilent Bioanalyzer to QC the libraries. The libraries were then pooled together to get the same numbers of reads from each cell when sequencing. Then we used the Illumina NextSeq 500 Sequencing System for high throughput sequencing that allows for 400M reads, permitting us to have broad coverage of each cell’s transcriptome. The Genome Analysis and Technology Core at UVA (RRID: SCR_018883) assisted in single cell barcoding, library construction, and sequencing. Raw reads were aligned to the mm39 reference genome and unique molecular identifier counting was done using the Cell Ranger pipeline (10X Genomics). Quality control and downstream analysis of the data was performed in R using Seurat v5^33^. Cells with mitochondrial gene expression <15%, hemoglobin gene expression < 0.075%, and gene counts between 200 and 5000 were retained for analysis. The UMI counts for the genes in each cell were log-normalized using the NormalizeData function in Seurat. The top 2000 highly variable features were identified, followed by integration and dimensionality reduction using the IntegrateData function and principal component analysis (PCA) in Seurat. The number of statistically significant principal components was determined using JackStraw scoring in Seurat, and these principal components were used for clustering with the Louvain algorithm to generate cell clusters (resolution = 0.5). To represent high dimensional data in two-dimensional space, we performed Uniform Manifold Approximation and Projection analysis using the PCA results. Doublets were detected and removed using the DoubletFinder package^53^, and then Louvain clustering and UMAP analysis were reperformed. A separate UMAP was generated for each group at each timepoint. The Louvain clusters were labeled/classified using canonical cell marker gene expression for each cluster. Clusters that had similar gene expression (*e*.*g*., fibroblasts, macrophages, etc.) were classified together. The FindMarkers function was used to identify significantly differentially expressed genes between MAP and NP implants.

### Cell-cell communication analysis

Cell-cell communication analysis was performed in CellChat v1.6^41^. The initial Seurat object was subsetted based on sample origin, and CellChat objects were created using the *createCellChat* function, and analysis was performed using the standard CellChat pipeline. The CellChat objects were then merged together using the *mergeCellChat* function. Plots were created using the *netVisual_aggregate, plotGeneExpression*, and *netVisual_bubble* functions.

### Processing cell lysate for ELISA and multiplex cytokine assay

After the hydrogel implants were digested as described above, a Cell Signaling Technologies cell lysis buffer (cat# 9803) was used to lyse the cells and isolate cytokines/proteins for analysis. The 1X buffer contained 20 mM Tris-HCL (pH 7.5), 150 mM NaCl, 1mM Na2EDTA, 1 mM EGTA, 1% Triton, 2.5 mM sodium pyrophosphate, 1 mM B-glycerophosphate, 1 mM Na3VO4, 1 µg/mL leupeptin, and 1 mM AEBSF. 0.5 mL of 1X cell lysis buffer was added to each sample and the solution was sonicated in a sonication water bath for 5 minutes. Samples were centrifuged at 18,000 x g for 10 minutes at 4°C and the supernatant from each tube was collected and diluted 1:5 in MilliQ water. For the multiplex cytokine assays, the cell lysate was submitted to the University Virginia Flow Cytometry core for analysis by a 32-plex cytokine multiplex assay (Luminex). The results of the cytokine multiplex assay were normalized to the total protein concentration within each implant via BCA assay. For the complement ELISAs, the cell lysate was serially diluted and ran with a C3a ELISA (Invitrogen) and C5a ELISA (Invitrogen) according to manufacturer protocols.

## RESOURCE AVAILABILITY

### Lead contact

Further information and requests for resources and reagents should be directed to and will be fulfilled by the lead contact, Don Griffin.

### Materials availability

This study did not generate new unique reagents.

### Data availability

- CyTOF and scRNA-seq data reported in this paper will be shared by the lead contact upon request.
- This paper does not report original code.
- Any additional information required to reanalyze the data reported in this paper is available from the lead contact upon request.

## ACKNOWLEDGMENTS

This work was funded by a NIBIB R21 Trailblazer [1R21EB028971-01A1], NIDDK R01 award [1R01DK126020], and an NIGMS R35 MIRA award [1R35GM160121]. C.A.R. was supported by the National Institute of General Medical Sciences of the National Institutes of Health under Award Number T32GM136615 and by the University of Virginia Dean’s Scholars Fellowship.

E.N. was supported by an R01 through the National Institute on Deafness and Other Communication Disorders [grant number 5R01DC020998–02]. J.O. was supported by the National Institute of General Medical Sciences of the National Institutes of Health under Award Number T32GM136615. J.M.S. was supported by the National Institutes of Health NHLBI under Award Number 1R01HL179312-01. The CyTOF and Luminex data for this manuscript was generated in the University of Virginia Flow Cytometry Core Facility (RRID:SCR_017829) and was partially supported by the NCI Grant (P30-CA044579). The authors acknowledge the University of Virginia Flow Cytometry Core for their assistance in conjugating some of the antibodies for CyTOF and for running of many the samples on the Helios cytometer for many hours. The authors acknowledge the University of Virginia Genome Analysis and Technology Core (RRID: SCR_018883) for generating the single-cell RNA sequencing data.

## AUTHOR CONTRIBUTIONS

C.A.R., E.N., J.O., R.H., J.S., D.A., and D.R.G. conceived the experiments. C.A.R. executed some animal experiments and the CyTOF experiments and analysis. E.N. performed some animal experiments, tissue histology staining and analysis, and the multiplex cytokine assays and analysis. J.O. and D.A. performed the RNAseq analysis. R.H. helped with the optimization of the CyTOF panel and protocols. C.A.R., E.N., J.O., R.H., J.S., D.A., and D.R.G. interpreted data and provided feedback. C.A.R. and D.R.G. wrote this manuscript and all authors contributed to editing of the manuscript. D.R.G. is the principal investigator.

## SUPPLEMENTAL INFORMATION

**Document S1. Figures S1-S2 and Table S1**

