## Supplemental text for "Microporous annealed particle scaffolds avoid foreign body response by down regulating complement-fibroblast-macrophage signaling loop"

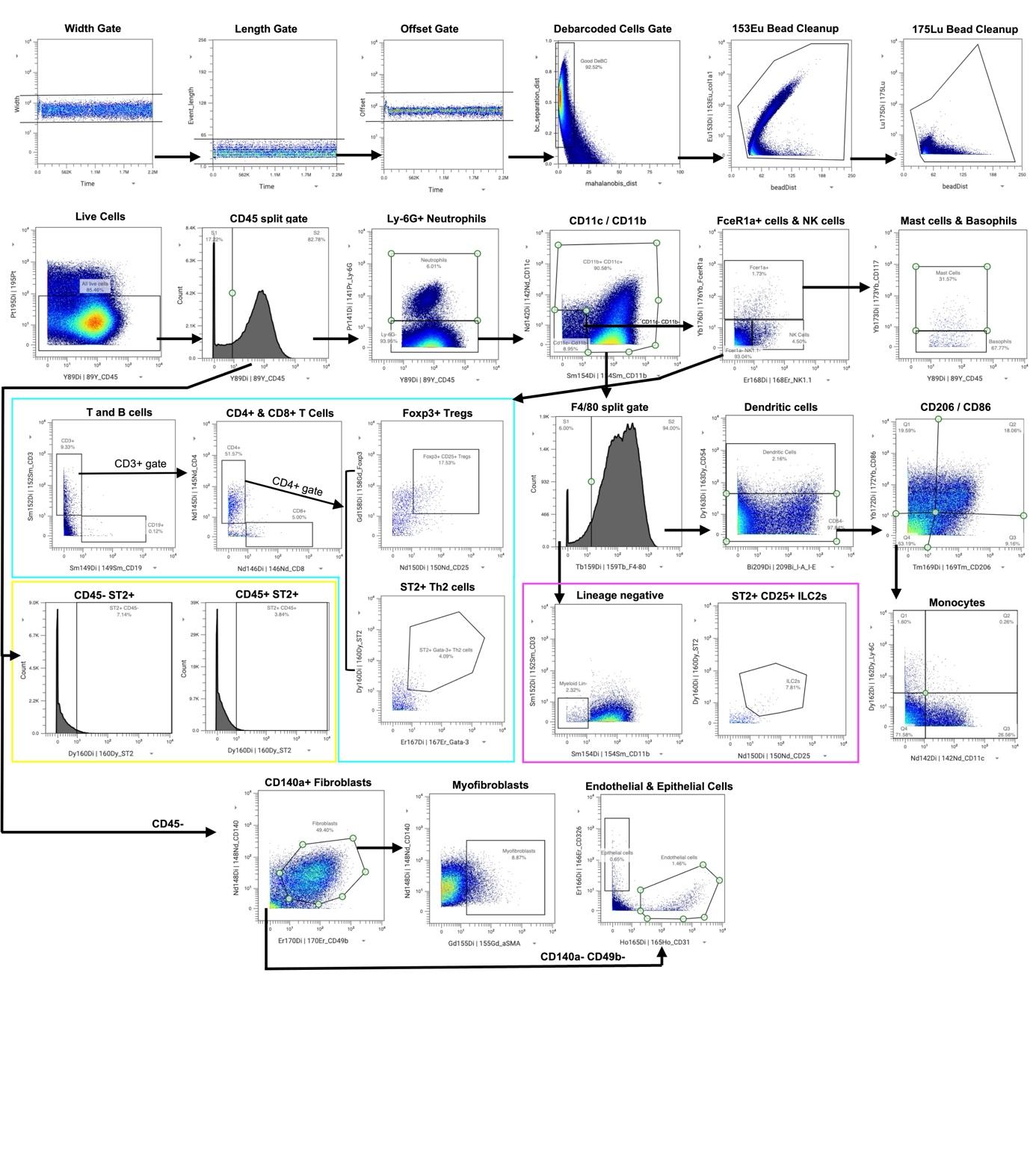

***Figure S1. Gating strategy for the CyTOF panel in OMIQ software.***

| **CyTOF Panel Antibodies** | | | |
| --- | --- | --- | --- |
| Isotope | Antibody | Clone | Dilution |
| Y89 | CD45 | 30-F11 | 0.25 |
| Pr141 | Ly-6G | rb6-8c5 | 0.125 |
| Nd142 | CD11c | n418 | 0.125 |
| Nd143 | CD69 | h1.2f3 | 2 |
| Nd145 | CD4 | rm4-5 | 0.25 |
| Nd146 | CD8a | 53-6.7 | 2 |
| Nd148 | CD140a (PDGFRa) | apa5 | 0.25 |
| Sm149 | CD19 | 6d5 | 0.5 |
| Nd150 | CD25 | 3c7 | 2 |
| Eu151 | CD140b (PDGFRb) | apb5 | 2 |
| Sm152 | CD3e | 145-2C11 | 2 |
| Eu153 | col1a1 | E8F4L | 0.03125 |
| Sm154 | CD11b | m1/70 | 0.0625 |
| Gd155 | aSMA | E8F4L | 0.03125 |
| Gd156 | CD90.2/Thy-1 | 30-H12 | 0.0625 |
| Gd158 | Foxp3 | FJK-16s | 1.7 |
| Tb159 | F4/80 | bm8 | 0.5 |
| Gd160 | ST2 | DIH9 | 2 |
| Dy161 | BAFF | polyclonal | 0.065 |
| Dy162 | Ly-6C | hk1.4 | 0.0625 |
| Dy163 | ICAM (CD54) | yn1/1.7.4 | 0.0625 |
| Dy164 | Ly-6A/E (sca-1) | d7 | 0.03125 |
| Ho165 | CD31 (PECAM) | 390 | 0.03125 |
| Er166 | CD326 (EpCAM) | g8.8 | 0.5 |
| Er167 | Gata-3 | TWAJ | 1.7 |
| Er168 | NK1.1 | PK136 | 2 |
| Tm169 | CD206 | C068C2 | 0.125 |
| Er170 | CD49b | HMa2 | 0.125 |
| Yb172 | CD86 | GL1 | 0.0625 |
| Yb173 | CD117 (ckit) | 2B8 | 0.125 |
| Yb174 | IL-17 | TC11-18H10.1 | 2 |
| Yb176 | FceR1a | MAR-1 | 2 |
| Bi209 | I-A/I-E | M5/114.15.2 | 0.125 |

***Table S1. List of CyTOF antibodies.*** Antibodies were purchased from Standard BioTools or conjugated to heavy metal isotopes in-house for this panel. The chosen dilutions for each antibody were optimized with multiple titration experiments (data not shown).

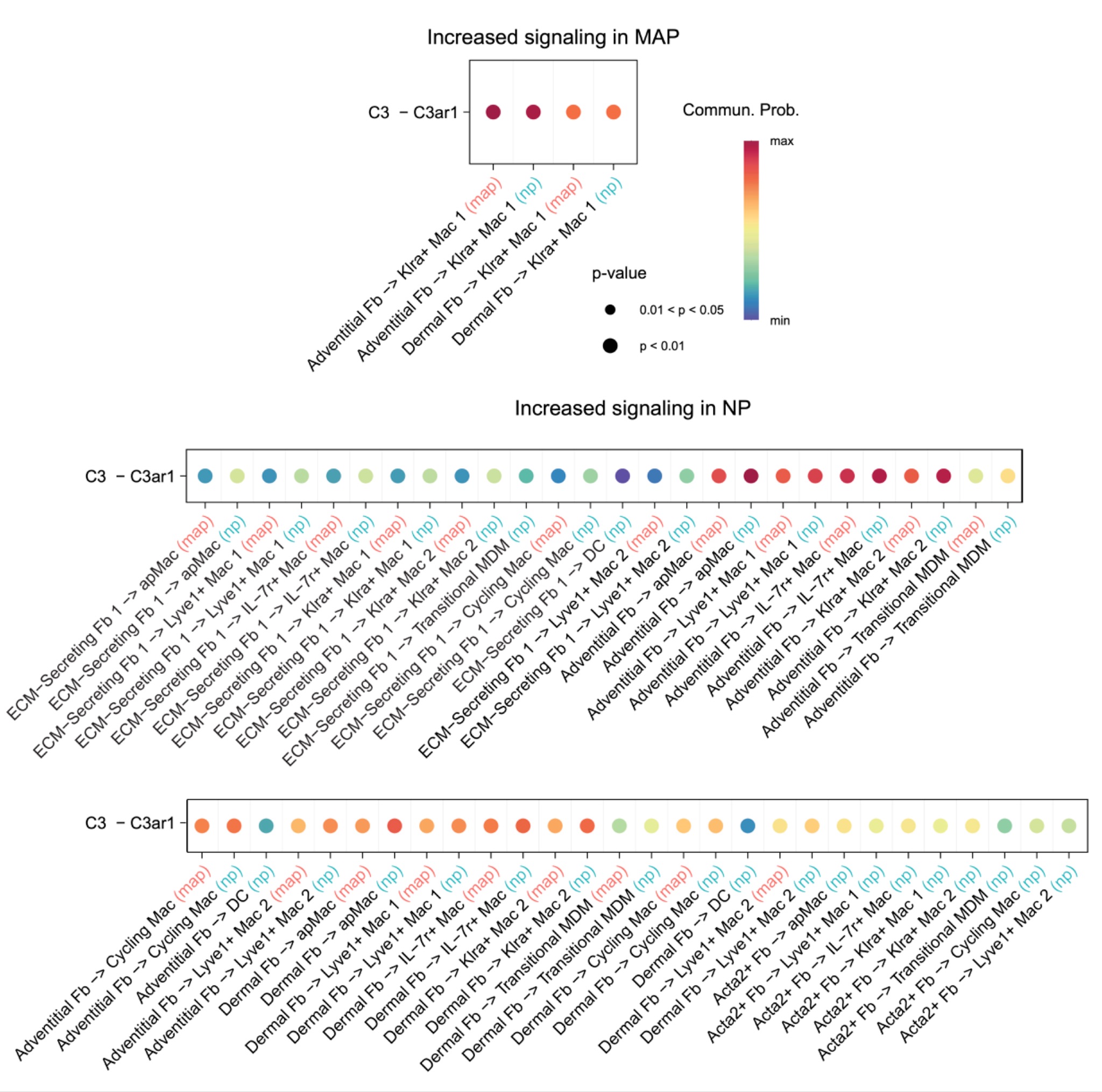

***Figure S2. CellChat analysis identifying complement signaling axis in MAP and NP implants***
